# Effect of Cysteine, Yeast Extract, pH Regulation and Gas Flow on Acetate and Ethanol Formation and Growth Profiles of *Clostridium ljungdahlii* Syngas Fermentation

**DOI:** 10.1101/2020.01.13.904292

**Authors:** Alba Infantes, Michaela Kugel, Anke Neumann

## Abstract

The fermentation of synthesis gas, or syngas, which consists mainly of CO, CO_2_ and H_2_ by acetogenic bacteria has the potential to help in transitioning from a fossil-fuel-based to a renewable bio-economy. *Clostridium ljungdahlii*, one of such microorganisms, has as main fermentation products acetate and ethanol. Multiple research efforts have been directed towards understanding how the metabolism and the product formation of this, and other acetogenic bacteria, can be directed towards increasing productivities and yields; nonetheless, transferring those findings to a particular set-up can prove challenging. This study used a well-established and robust fed-batch fermentation system with *C. ljungdahlii* to look into the effects of different fermentation pH profiles, gas flow, and the supplementation with additional yeast extract or cysteine on growth, product formation ratios, yields, and productivities, as well as gas consumption. Neither yeast extract nor cysteine supplementation had a noticeable impact on cell growth, product formation or overall gas consumption. The lowering of the pH proved mainly detrimental, with decreased productivities and no improvement in ethanol ratios. The most notable shift towards ethanol was achieved by the combination of lowering both the pH and the gas flow after 24 h, but with the caveat of lower productivity. The obtained results, unexpected to some extent, highlight the necessity for a better understanding of the physiology and the metabolic regulation of acetogenic bacteria in order for this process to become more industrially relevant.

## INTRODUCTION

In the current scenario of a growing world population and decreasing resources, together with the environmental implications of fossil fuel combustion, alternatives sources for fuels and chemicals need to be found. The dependence of our society on fossil fuels is clear: a vast majority of everyday materials, as well as primary energy, are derived from fossil fuels (Edenhofer et al., 2014). In contrast, the fermentation of synthesis gas (syngas) by acetogenic bacteria can provide with an environmentally-friendly and renewable alternative for the production of low-carbon fuels and chemicals, and is receiving ever more attention (Hu et al., 2016; Liew et al., 2013; Richter et al., 2016).

Syngas consists of a mixture of mainly CO, H_2_ and CO_2_, and is derived from the gasification of biomass. This is advantageous in comparison to the fermentation of biomass-derived sugar feedstocks since the lignin fraction becomes accessible (Liew et al., 2016). Carboxydotrophic and homoacetogenic bacteria such as *Clostridium ljungdahlii* can grow by using the carbon and electrons derived from syngas, thanks to their unique carbon-fixating Wood-Ljundahl pathway. Their primary end-product is acetic acid and ethanol, but other products like butyrate or butanol have also been described (Bengelsdorf et al., 2018; Henstra et al., 2007). Given the fact that these microorganisms are becoming more relevant, the understanding of the fermentation process and their product formation profile is of valuable interest.

Acetogenic bacteria growing on CO_2_/H_2_ and/or CO experiment a metabolic shift similar to that seen in ABE Clostridia (Liew et al., 2013), which display a biphasic fermentation profile: during exponential growth, carboxylic acids are produced together with H_2_ and CO_2_ (acidogenic phase), while during the stationary phase these acids are taken up and solvents are formed (solventogenic phase). This metabolic change is accompanied by a significant change in gene expression (Lee et al., 2008). Despite the strategy similarities of both ABE and syngas-fermenting Clostridia, no gene expression regulation could be found as the mechanism driving the shift in the syngas-fermenting microorganism (Richter et al., 2016). Physiologically, temperature, pH, acetic acid concentration, and nutrient limitation are regarded as factors that can induce a transition to non-growth conditions and solventogenesis (Liew et al., 2013; Mohammadi et al., 2011).

By increasing or decreasing certain medium components, biomass, gas consumption and product formation can be affected. For instance, Abubackar et al. (2012) showed that increasing the cysteine concentration in the medium enhanced the ethanol production of *Clostridium autoethanogenum*. The work of Saxena and Tanner (2012), showed that yeast extract and trace metals were required for *Clostridium ragsdalei* to grow.

Moreover, based on a proteome analysis, Richter et al. (2016) found that the genes for the sulfate reduction in the sulfur-assimilation pathway in *C. ljungdahlii* are absent. Therefore, they suggested replacing the sulfate that is usually present in the syngas fermentation medium by sulfide or cysteine. In this context, a medium containing no sulfate, but cysteine as the sulfur source has already been reported to support growth for *Clostridium autoethanogenum*, *Clostridium ljungdahlii* and *Clostridium ragsdalei* (Annan et al., 2019).

Yeast extract is a crucial part of the medium since its removal does not support the growth of syngas-fermenting microorganisms (Barik et al., 1988). Even so, the reduction of its concentration in the medium has been shown to be beneficial in terms of ethanol production (Abubackar et al., 2012; Barik et al., 1988; Phillips et al., 1993; Vega et al., 1989). Still, the necessity for complex and not well understood medium components in the syngas fermentation medium is remarkable: one study showed that when substituting yeast extract with CSL (corn steep liquor), the supplementation with 0.5 g/L of yeast extract or trace metals, NH_4_^+^ and cysteine was necessary to support growth of *C. ragsdalei.* Even CSL itself could not be reduced below 10 g/L without experimenting a reduction in ethanol production (Saxena and Tanner, 2012). Cotter et al. (2009a) assessed the effect of nitrogen limitation to achieve stable resting cultures for the production of ethanol. When they removed nitrogen from the media, neither *C. ljungdahlii* nor *C. autoethanogenum* were able to maintain a stable cell density. They concluded that even non-growing cultures of *C. ljungdahlii* and *C. autoethanogenum* required organic nitrogen sources to prevent decaying cell densities.

On the other hand, providing additional nutrients in the form of yeast extract to supports cell growth had a positive effect on acetic acid production in a study by Barik et al. (1988). This agrees with acetate being a growth-related product (Barik et al., 1988; Richter et al., 2016), since higher biomass resulted in an enhanced production.

Certainly, not only nutrients affect the outcome; pH also plays an essential role in the fermentation performance: it significantly impacts the behavior of the microorganism, affecting both growth rate and product formation. A drop in the external pH might be a way to induce the production of more reduced compounds, such as ethanol (Abubackar et al., 2012; Barik et al., 1988; Phillips et al., 1993). For instance, a lower fermentation medium pH resulted in a switch towards ethanol production in *C. carboxidivorans* and *Butyribacterium methylotrophicum* (Ahmed et al., 2006; Worden et al., 1991). In a separate study with *B. methylotrophicum*, the pH was lowered from 6.8 to 6.0 only after reaching the stationary phase, which caused an increase in butyrate production (Diender et al., 2016). Similarly, Klasson et al. (1993) firstly grew a dense *C. ljungdahlii* culture at a pH of 5.0, and only then was the pH lowered between 4.0 and 4.5 to enhance ethanol production. Even so, Kundiyana et al. (2011) reported that lowering the pH below 6.0 did not produce a beneficial effect on ethanol production on *C. ragsdalei*. It must also be taken into account that acetate in its undissociated form is lipophilic and freely diffuses through the cell membrane, which results in the move of H^+^ across the transmembrane gradient, lowering the intracellular pH (Kundiyana et al., 2011). If the pH drops too low, it might negatively impact the culture since the microorganism could struggle to maintain a neutral intracellular pH (Cotter et al., 2009a; De Tissera et al., 2017; Mohammadi et al., 2011).

Another point of focus of this study was the gas consumption profile of the culture. Previous fed-batch fermentations performed in our group (data not shown) have shown that after achieving and maintaining complete gas usage, CO_2_ and H_2_ consumption stops consistently after ~60 h, eventually followed by the halt of CO consumption. CO is consumed completely, and therefore not accumulating in the medium for at least 48 h before this effect could be observed, so the idea that this could be caused by CO inhibition was discarded. Besides, since a halt in biomass growth accompanies this behavior, we hypothesized that our medium might be deficient in a main nutrient.

Taking all this into account, this study focused on assessing the effect of nutrients - yeast extract and sulfur, in the form of cysteine - on the fermentation of syngas by *C. ljungdahlii*, in terms of products, biomass formation, and gas consumption. The impact of pH in the acetate/ethanol ratio, as well as the influence of the amount of substrate fed in the form of the gas flow rate, were investigated.

## METHODS

### Microorganism and Medium

*C. ljungdahlii* DSM 13528 was used to perform the fermentations. Unless otherwise stated, all chemicals were acquired from Carl Roth GmbH + Co. KG (Germany) or Sigma-Aldrich Inc. (Germany).

Both the pre-cultures and the fermentation media was based on Tanner (2007). It contained: 20 g/L 2-(*N*-morpholino) ethansulfonic acid (MES) (20), 0.5 g/L yeast extract (BD, USA), 2 g/L NaCl, 2.5 g/L NH_4_Cl, 0.25 g/L KCl, 0.25 g/L KH_2_PO_4_, 0.5 g/L MgSO_4_•7 H_2_O, 0.1 g/L CaCl_2_•2 H_2_O, 10 mL trace element solution, 10 mL vitamin solution and 0.001 g/L resazurin. The pH was adjusted to 5.9 using KOH before autoclaving at 121 °C. After that, 0.6 g/L Cysteine-HCl•H_2_O were added to each fermenter, or 1 g/L to each serum flask in the case of the pre-culture. Trace element solution contained: 2 g/L nitrilotriacetic acid, 1 g/L MnSO_4_•H_2_O, 0.567 g/L FeSO_4_•7 H_2_O, 0.2 g/L CoCl_2_•6 H_2_O (Riedel-de Haën, Germany), 0.2 g/L ZnSO_4_•7 H_2_O, 0.02 g/L CuCl_2_•2 H_2_O, 0.02 g/L NiCl_2_•6 H_2_O, 0.02 g/L Na_2_MoO_4_•2 H_2_O, 0.02 g/L Na_2_SeO_3_•5 H_2_O and 0.022 g/L Na_2_WO_4_•2 H_2_O. Vitamin solution contained: 2 mg/L biotin, 2 mg/L folic acid, 10 mg/L pyridoxine (Alfa Aesar, Germany), 5 m g/L thiamine-HCl, 5 mg/L riboflavin, 5 mg/L niacin, 5 mg/L calcium-pantothenate, 5 mg/L cobalamin, 5 mg/L 4-aminobenzoic acid, and 5 mg/L lipoic acid (Cayman Chemical, USA).

The pre-cultures for each experiment were freshly prepared, starting from a single glycerol stock. Glycerol stocks were produced from a 48 h grown culture. 5 mL of the culture was aseptically and anaerobically removed and dispensed into a sterile, anaerobised Hungate-type culture tube. The culture was then centrifuged for 5 minutes at 4°C and 3000 g. The supernatant was then discarded, and the pellet was resuspended in 1 mL of anaerobic and sterile freezing solution, made with equal volumes of culture media and a 50 vol-% glycerol solution.

For the pre-culture, a glycerol stock frozen at −80 °C was thawed and its entire volume was anaerobically and sterilely dispensed into an anaerobic serum flask containing 50 mL of the Tanner medium. The carbon source used for the pre-cultures was 10 g/L of fructose. This culture was allowed to grow for 48 h at 37 °C without shaking. Two subsequent passages with the same cultivation conditions were performed, where a 10 % inoculum volume was added to serum flasks containing fresh medium. The fermenters were inoculated with a 10 % inoculum volume (150 mL) and each one was inoculated from an individual serum flask.

### Fermentation conditions

All fermentations were carried out in Minifors® bench-top stirred tank reactors (Infors-HT, Switzerland), which have a total volume of 2.5 L. The working liquid volume was 1.5 L. All experiments were performed in triplicates.

The gas for the fermentation was supplied with a microsparger, while the gas flow rate was controlled via a mass flow controller (MFC) red-y smart series, from Vögtlin Instruments (Switzerland).

The temperature of the fermenter was kept at 37 °C, pH was controlled at 5.9 with 4 M KOH, and stirring was regulated at 800 rpm.

Anaerobic conditions were ensured after autoclaving by sparging the fermenters with N_2_ for 2 h. Following this, the gas supply was changed to syngas with a flow rate of 50 mL/min for at least 3 h until just before inoculation, when the gas flow rate was adjusted as required.

A detailed description of the fermenter set-up can be found in Oswald et al. (2016). The gas flow rate being fed into the fermenters was controlled at 18 mL/min. For all the fermentations, a pure gas mixture was used, with the following composition: 32.50 vol-% CO, 16.00 vol-% CO_2_, 32.50 vol-% H_2_ and 19 vol-% N_2_ (Air Liquide, France).

### Experimental set-up

Six experiments were conducted, where the effect of different gas flows, pH, initial yeast extract concentration and initial cysteine concentration were observed. Each experiment was performed as a triplicate (unless otherwise stated), and all fermentations were carried out for approximately 93 h. A detailed description of each set-up can be found in Table 1.

**Table 1.** Experimental set-up. All fermentations were done as triplicates (n=3), except for set-up 3, where one fermenter was kept unaltered (3a), and two fermenters were treated (3b).

| Set-up | Medium | pH | Gas flow |
| --- | --- | --- | --- |
| 1 (Standard) | Standard | 5.9 | 18 mL/min |
| 2 | Standard | No pH regulation | 18 mL/min |
| 3a | Increased cysteine to 1 g/L | 5.9 | 18 mL/min |
| 3b | Increased cysteine to 1 g/L | After 68 h, pH changed to 5.4 | 18 mL/min |
| 4 | Doubled yeast extract concentration to 1 g/L | 5.9 | 18 mL/min |
| 5 | Standard | 24 h at pH 5.9, then let drop to pH 4.8 and hold | 18 mL/min |
| 6 | Standard | 24 h at pH 5.9, then let drop to pH 4.8 and hold | 24 h at 18 mL/min, then decreased to 12.6 mL/min |

### Analytical Methods

The fermenters’ off-gas were analyzed using a GC-2010 Plus AT gas chromatograph (GC) (Shimadzu, Japan), with a ShinCarbon ST 80/100 Column (2 m × 0.53 mm ID, Restek, Germany) and an Rtx-1 capillary column (1 μm, 30 m × 0.25 mm ID, Restek, Germany). The detector used was a thermal conductivity detector with helium as carrier gas. The column flow rate was 3 mL/min, with an oven temperature of 40 °C for 3 min followed by a ramp of 35 °C/min. The total analysis time was 7.5 min. Data obtained was subsequently evaluated as described in (Oswald et al., 2016).

The sampling regime was as follows: four samples of 2 mL were taken daily at 2 – 3 h intervals, with no sample collection taking place overnight. These were then used for OD (optical density) determination and left-over fructose and products (acetate and ethanol) concentration. OD (optical density) was determined at 600 nm.The sample collection, its treatment, and off-line analysis are described in detail in Oswald et al. (2016).

The OD (optical density) and cell dry weight (CDW) correlation was determined as the average of 12 fermentations under comparable conditions (data not shown), with a resulting factor of CDW/OD = 0.30 ± 0.04 g/L ⋅ OD.

### Calculation of product formation parameters using different metrics and at specific phases of the fermentation

First of all, the terminology used here is clarified. “Substrate” refers to the syngas fed into the bioreactor, that is, CO, CO_2_ and H_2_.

The amount of substrate “fed” corresponds to the total amount of CO, CO_2_ and H_2_, in mol, that has been sparged into the bioreactor during a specific time interval.

The term “usage” refers to the difference of a particular gas species between the amount fed into the reactor and the amount measured in the off-gas stream (in mol). A negative number indicates production.

Carbon “fixation” refers to the amount of CO_2_ or CO, in mol, that got incorporated into products or biomass.

For an accurate analysis, in the case of CO, it is necessary to distinguish between CO used and CO fixed. “CO fixed” refers specifically to the amount of this substance assimilated by the cells to products or biomass. “CO used” comprises both the CO fixed and the amount of CO that is converted to CO_2_ by the bacteria which does not get incorporated and is released with the out-gas.

Hence, in the absence of any other carbon source, if the amount of CO_2_ in the off-gas is larger than the amount being fed (i.e., the CO_2_ usage value is positive), it is an indication of CO being converted to CO_2_. This amount of produced CO_2_ from CO (in mol) must be subtracted from the “used” amount of CO (in mol) to obtain the actual amount of CO fixed.

Taking this into account, two scenarios are possible: Firstly, if CO_2_ is indeed produced from CO, the value of the (perceived) CO usage will be higher than the actual amount of CO fixed into products and biomass. Secondly, if no CO_2_ is produced,, then the amount of CO fixed is equal to the amount of CO used. For clarity, a short overview of the calculation is given below:

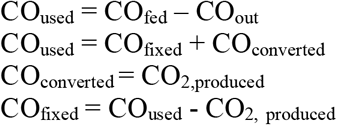

Regarding yield calculations, they are all given here in gram of product (the sum of acetate and ethanol, in grams) per gram of substrate (the sum of CO_2_, CO and H_2_, in grams). Three approaches were used: yield per substrate fed (Y_P/S, fed_), yield per substrate used (Y_P/S, used_), and yield per substrate fixed (Y_P/S, fixed_). In the latter case, this includes the amount of CO fixed, the amount of H_2_, and, if any, the amount of CO_2_ used.

In order to be able to analyze and compare the data between both fermentations, the product formation parameters yield and productivity, as well as the acetate to ethanol ratio were calculated for the complete run, and up to the point when maximum CO consumption stopped (Tables 3 and 4).

Endpoint calculations were done using the values measured with the sample taken immediately before terminating the fermentation.

The maximum CO fixation interval was determined by identifying the period where the CO fixation reached a value of 85 % or higher. Calculations were done from the starting of the fermentation to the last point when CO fixation was above 85 %. Due to limitations in the amount of samples that could be withdrawn, the measurements from the sample closest to that point are used.

The interval of maximum overall usage is determined according to the gas consumption profile. The sum of CO fixation and CO_2_ and H_2_ usage for each measured point is calculated throughout the fermentation; note that only if no CO_2_ is produced then CO used equals CO fixed. The maximum value achieved is defined as the maximum overall usage. The interval of maximum overall usage is the period during which the sum of the usage value of the three gaseous substrates is ≥ 85 % of the mentioned maximum.

## RESULTS

If not otherwise stated, all fermentations were done as triplicates (n=3), and the results are presented here as the average.

### Effect of medium components

The standard fermentation (set-up 1) achieved a final acetate and ethanol concentration of 20.1 g/L and 2.0 g/L, respectively, after 95 h (Figure 1A). At 69 h, 15.0 g/L of acetate and 0.9 g/L of ethanol had been formed. In the case of the increased cysteine, at 68 h, 13.6 g/L of acetate and 0.9 g/L of ethanol had been formed. After 95 h, the concentration of products in the reactor kept at pH 5.9 was 16.6 g/L of acetate and 2.0 g/L of ethanol. During the fermentation with 1 g/L of cysteine (set-up 3), it became clear that the behavior of the culture was equivalent to that of the one performed with standard medium (set-up 1). In two fermenters (set-up 3b) the pH was lowered after 68 h (figures 1B and 3B), but one fermenter (set-up 3a) was kept at 5.9 to corroborate that an increased cysteine concentration did also not affect the behavior of the microorganism later in the run (Figure 1C).

**Figure 1:**
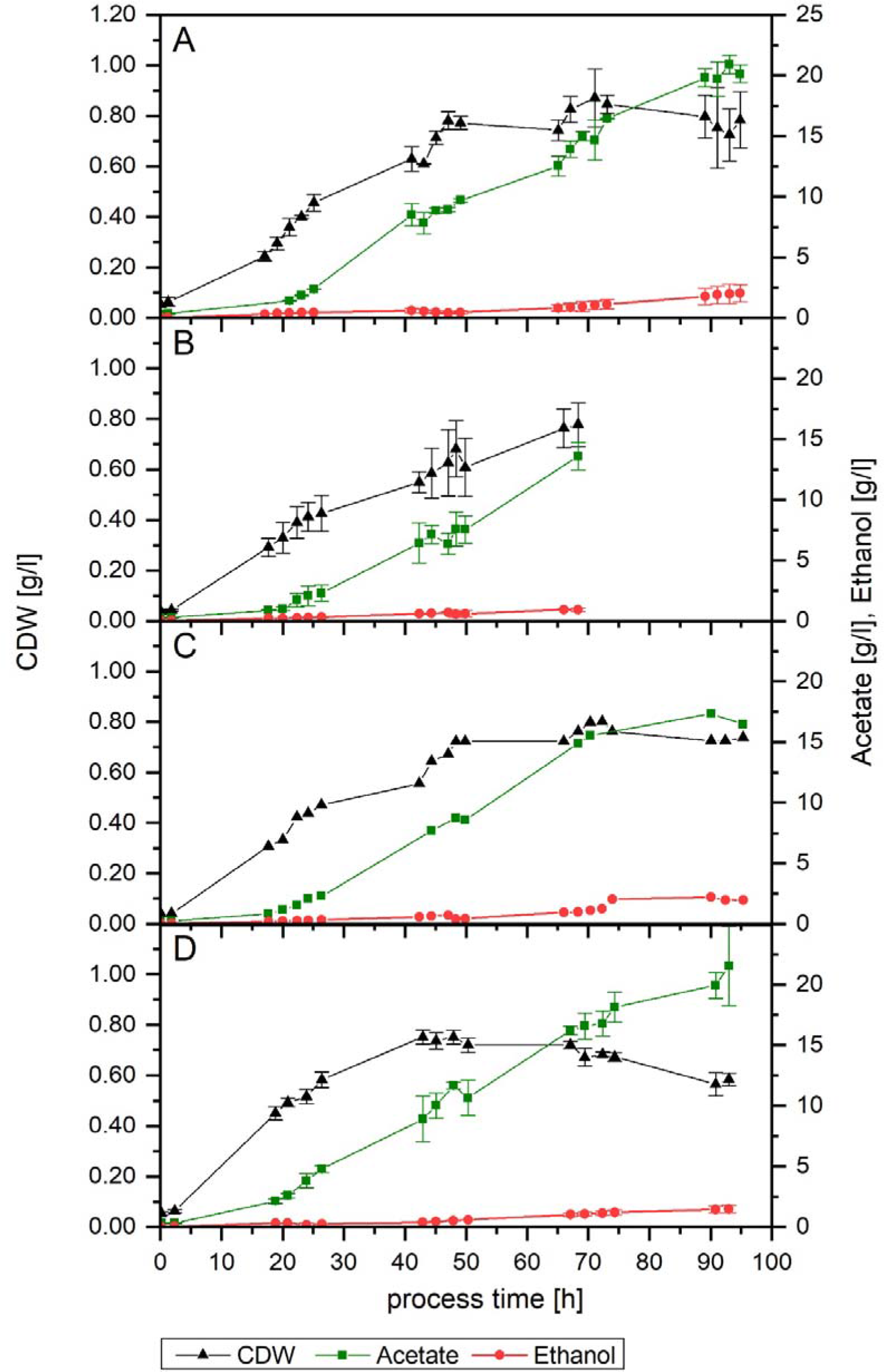
Growth and product formation of the standard set-up 1 (A), set-up 3 with increased cysteine (B and C) and set-up 4 with increased yeast extract (D). Figure B shows set-up 3b (up to 68 h, point where pH was changed). Figure C shows set-up 3a. Average values of the triplicates (n=3) for cell dry weight (CDW, black triangles), acetate (green squares) and ethanol (red dots), except for Figure C where only 1 fermented is depicted.

Both set-ups 1, 3a, and 3b followed a very similar growth pattern up to 68 h. The standard set-up 1 reached a final and a maximum cell dry weight (CDW) of 0.8 g/L and 0.9 g/L, respectively. In this fermenter, at 67 h, the CDW concentration was 0.8 g/L. In the case of increased cysteine, the same value was achieved at 68 h. For the fermenter left unaltered, set-up 3a, the CDW at 95 h of process-time and the maximum value reached were 0.7 and 0.8 g/L, respectively.

Concerning the influence of an increased yeast extract concentration (set-up 4), a comparable final amount of acetate was formed (21.5 g/L), but only 1.4 g/L of ethanol was produced (Figure 1D). In terms of biomass, the final reached value was lower (0.6 g/L), as well as its maximum (0.8 g/L at 48 h), resulting in a notably higher Y_P/X_ value: 41 g/g compared to 29 g/g in both set-ups 1 and 3a.

Substrate consumption graphs are depicted in detail in Figure 2; because of the difference of the gas consumption profiles between both fermenters in set-up 3b, only fermenter “c” was included in the averages in this figure. Both medium modifications performed similarly to the standard set-up 1 as to the duration of the maximum overall usage of the substrate, but the starting and ending time did differ, with set-up 4 (increased yeast extract) starting earlier. When looking solely at CO fixation, set-ups 3a, 3b, and 4 behaved alike, with the maximum CO fixation lasting around 10 h less compared to the standard conditions (Table 2). The off-gas profile for set-ups 1, 3 and 4 are shown in Supplementary Figure 1.

**Table 2.**
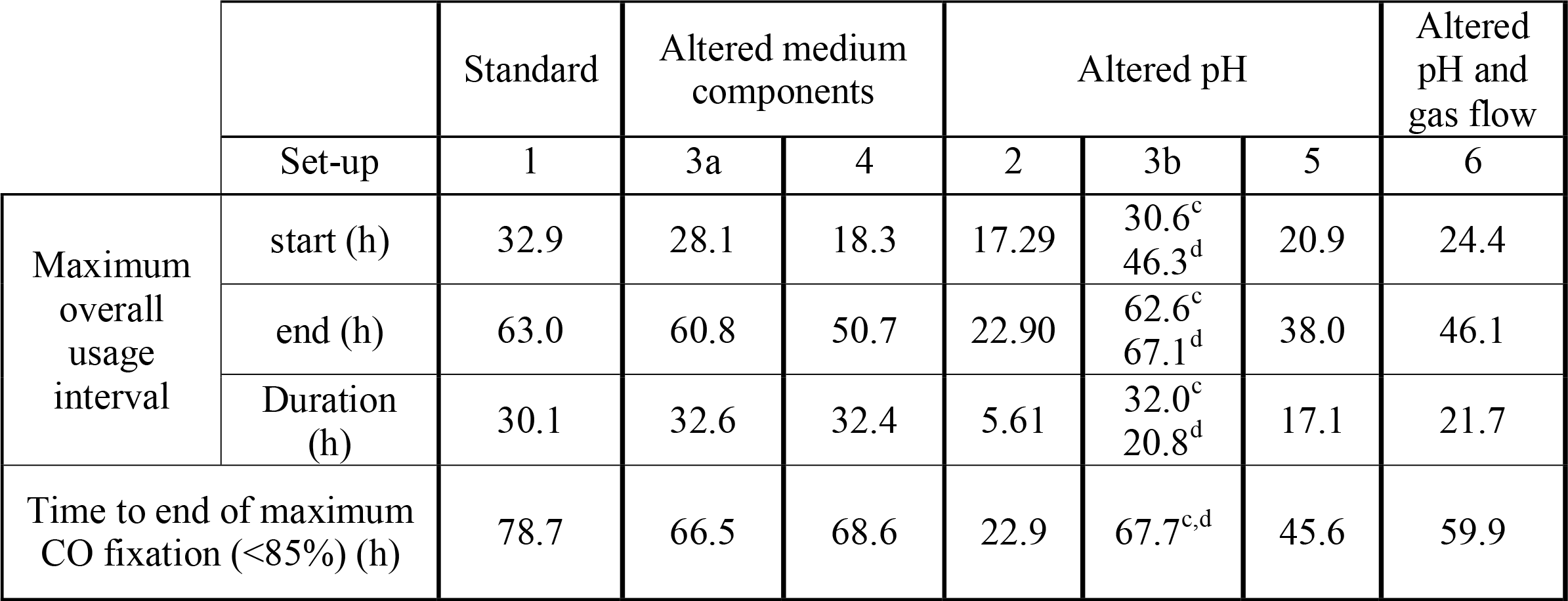
Gas consumption profiles. All values given as an average of a triplicate (n=3), except for the following: 3a - values of the fermenter where pH was not altered; 3b - average of the two fermenters where the pH was changed after 68 h to 5.4, with superscripts c and d designating the value for each individual fermenter, due to the divergence observed.

|  |  | Standard | Altered medium components |  | Altered pH |  |  | Altered pH and gas flow |
| --- | --- | --- | --- | --- | --- | --- | --- | --- |
|  | Set-up | 1 | 3a | 4 | 2 | 3b | 5 | 6 |
| Maximum overall usage interval | start (h) | 32.9 | 28.1 | 18.3 | 17.29 | 30.6<sup>c</sup><br>46.3<sup>d</sup> | 20.9 | 24.4 |
|  | end (h) | 63.0 | 60.8 | 50.7 | 22.90 | 62.6<sup>c</sup><br>67.1<sup>d</sup> | 38.0 | 46.1 |
|  | Duration (h) | 30.1 | 32.6 | 32.4 | 5.61 | 32.0<sup>c</sup><br>20.8<sup>d</sup> | 17.1 | 21.7 |
| Time to end of maximum CO fixation (<85%) (h) |  | 78.7 | 66.5 | 68.6 | 22.9 | 67.7<sup>c,d</sup> | 45.6 | 59.9 |

**Figure 2:**
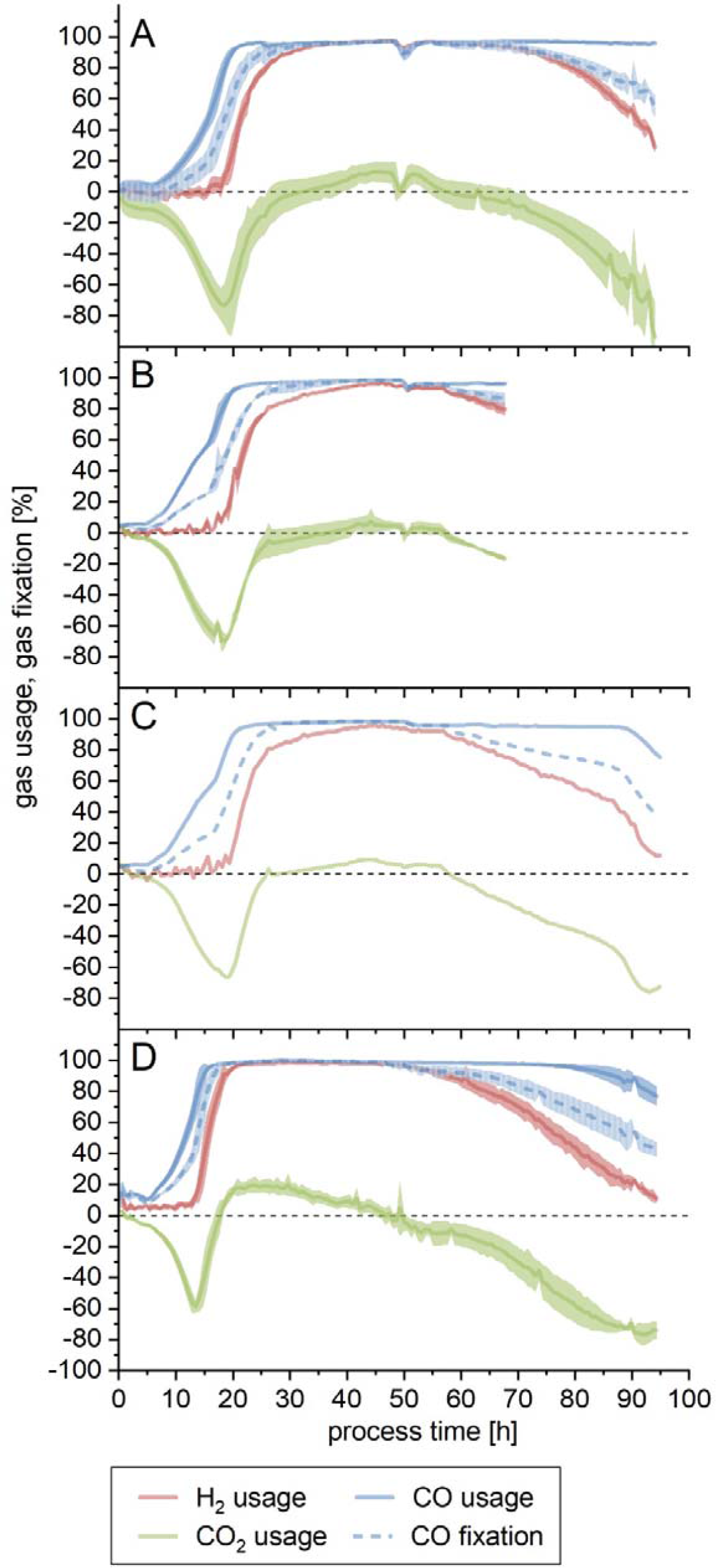
Substrate usage or fixation for the standard set-up 1 (A), set-up 3 with increased cysteine (B and C) and set-up 4 with increased yeast extract (D). Figure B shows set-up 3b up to 68 h (point where pH was changed, not shown here). One fermenter has been left out of the averages due to being remarkably delayed in comparison with the other two. Figure C shows set-up 3a, where one fermenter was kept at pH 5.9. Usage is for H_2_ (red line), CO_2_ (green line) and CO (blue line). CO fixation is depicted by the dotted blue line. The calculated difference between amount of substance flow rate fed into the bioreactor and the amount of substance flow rate detected in the off-gas is shown here as a percentage. For the CO fixation, if the CO_2_ usage was negative, the amount of CO_2_ produced was subtracted from the amount of (perceived) CO used. Except where otherwise stated, lines show the average of a triplicate (n=3), while the lighter colored areas depict the standard deviation.

Concerning the overall yields (calculated up to the end of the fermentation), the most significant difference is the Y_P/X_ in set-up 4, as mentioned above (Table 3). Moreover, this fourth experiment had the highest productivity among all fermentations, despite the reduction in the amount of ethanol produced.

**Table 3.**
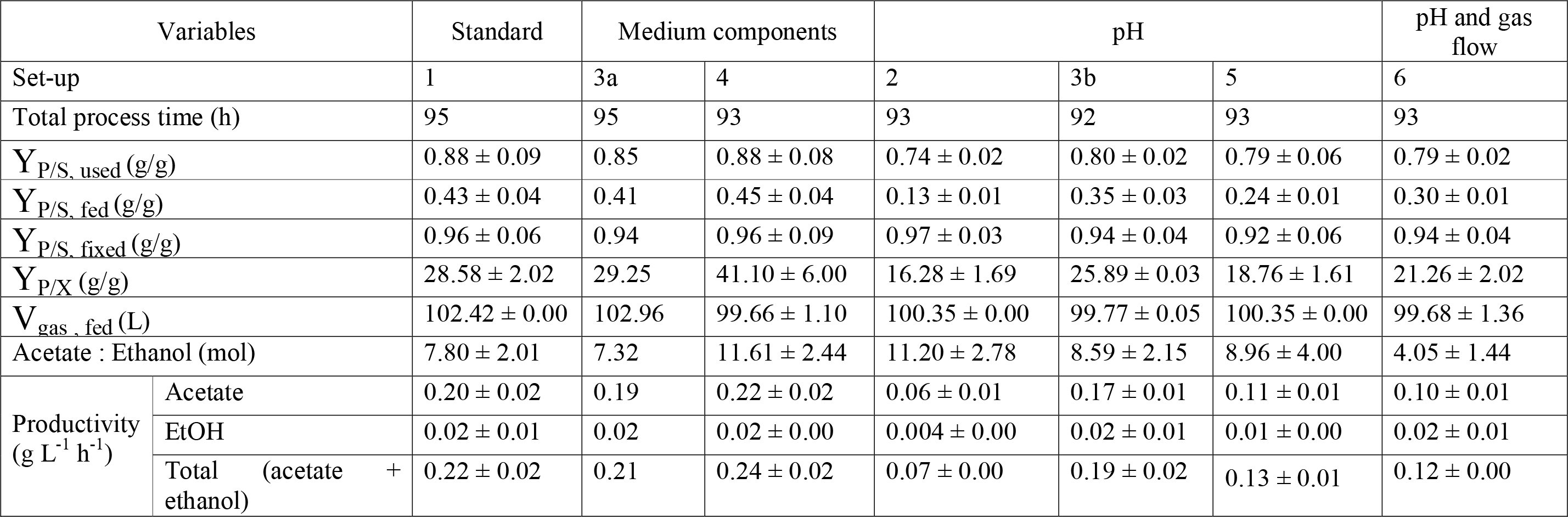
Fermentation outcomes, yields, and productivities for the complete run. 3a: calculated for the fermenter where pH was not altered. 3b: referring to the pair of fermenters where the pH was changed after 68 h to 5.4. Y_P/S_, (g/g) = gram of products (acetate and ethanol) formed per gram of substrate (CO, CO_2_ and H_2_). This has been calculated per grams of substrate fed, used and fixed. Y_P/X_ (g/g) = gram of product (acetate and ethanol) per gram of biomass (cell dry weight). Values are given as the average of a triplicate (n=3) with standard deviations, except for set-up 3a, where only the values for the fermenter left unaltered at pH 5.9 are shown, and set-up 3b, where the average of the two fermenters on which pH was modified is given.

| Variables |  | Standard | Medium components |  | pH |  |  | pH and gas flow |
| --- | --- | --- | --- | --- | --- | --- | --- | --- |
| Set-up |  | 1 | 3a | 4 | 2 | 3b | 5 | 6 |
| Total process time (h) |  | 95 | 95 | 93 | 93 | 92 | 93 | 93 |
| $Y_{P/S, \text{ used}}$ (g/g) | | $0.88 \pm 0.09$ | 0.85 | $0.88 \pm 0.08$ | $0.74 \pm 0.02$ | $0.80 \pm 0.02$ | $0.79 \pm 0.06$ | $0.79 \pm 0.02$ |
| $Y_{P/S, \text{ fed}}$ (g/g) | | $0.43 \pm 0.04$ | 0.41 | $0.45 \pm 0.04$ | $0.13 \pm 0.01$ | $0.35 \pm 0.03$ | $0.24 \pm 0.01$ | $0.30 \pm 0.01$ |
| $Y_{P/S, \text{ fixed}}$ (g/g) | | $0.96 \pm 0.06$ | 0.94 | $0.96 \pm 0.09$ | $0.97 \pm 0.03$ | $0.94 \pm 0.04$ | $0.92 \pm 0.06$ | $0.94 \pm 0.04$ |
| $Y_{P/X}$ (g/g) | | $28.58 \pm 2.02$ | 29.25 | $41.10 \pm 6.00$ | $16.28 \pm 1.69$ | $25.89 \pm 0.03$ | $18.76 \pm 1.61$ | $21.26 \pm 2.02$ |
| $V_{\text{gas, fed}}$ (L) | | $102.42 \pm 0.00$ | 102.96 | $99.66 \pm 1.10$ | $100.35 \pm 0.00$ | $99.77 \pm 0.05$ | $100.35 \pm 0.00$ | $99.68 \pm 1.36$ |
| Acetate : Ethanol (mol) | | $7.80 \pm 2.01$ | 7.32 | $11.61 \pm 2.44$ | $11.20 \pm 2.78$ | $8.59 \pm 2.15$ | $8.96 \pm 4.00$ | $4.05 \pm 1.44$ |
| Productivity<br>(g L<sup>-1</sup> h<sup>-1</sup>) | Acetate | $0.20 \pm 0.02$ | 0.19 | $0.22 \pm 0.02$ | $0.06 \pm 0.01$ | $0.17 \pm 0.01$ | $0.11 \pm 0.01$ | $0.10 \pm 0.01$ |
| | EtOH | $0.02 \pm 0.01$ | 0.02 | $0.02 \pm 0.00$ | $0.004 \pm 0.00$ | $0.02 \pm 0.01$ | $0.01 \pm 0.00$ | $0.02 \pm 0.01$ |
| | Total (acetate + ethanol) | $0.22 \pm 0.02$ | 0.21 | $0.24 \pm 0.02$ | $0.07 \pm 0.00$ | $0.19 \pm 0.02$ | $0.13 \pm 0.01$ | $0.12 \pm 0.00$ |

For easier comparison, since all fermentations where run for approximately 93 h, but each stopped consuming the gaseous substrates at different times, yields and productivities were also calculated up to the point when maximum CO fixation came to an end, as found in Table 4. Set-ups 1, 3a, 3b, and 4 performed likewise when compared up to the point when maximum CO fixation stopped. The most noticeable difference is the lower Y_P/S, fed_ achieved by set-up 3b. The highest converted amount of carbon fed into products (Y_P/S, fed_) was reached by the standard fermentation (0.51), while set-up 3b was the lowest (0.38). Nonetheless, the latter achieved a comparable yield of products per carbon fixed (Y_P/S, fixed_). In terms of gram of product formed per gram of biomass (Y_P/X_), the difference seen on the end-of-process yields is already to be found here, with set-up 4 achieving the highest value. The acetate to ethanol ratio also differs slightly during this phase, with the most acetate per mol of ethanol being produced by the standard fermentation, contrasting with the results seen when looking at the end-of-process values.

**Table 4.** Fermentation outcomes, yields, and productivities calculated up to the point when maximum CO fixation stopped. 3a: calculated for the fermenter where pH was not altered. 3b: referring to the pair of fermenters where the pH was changed after 68 h to 5.4. Y_P/S,_ (g/g) = gram of products (acetate and ethanol) formed per gram of substrate (CO, CO_2_ and H_2_). This has been calculated per grams of substrate fed, used and fixed. Y_P/X_ (g/g) = gram of product (acetate and ethanol) per gram of biomass (cell dry weight). Values are given as the average of a triplicate (n=3) with standard deviations, except for set-up 3a, where only the values for the fermenter left unaltered at pH 5.9 are shown, and set-up 3b, where the average of the two fermenters on which pH was modified is given.

| Variables |  | Standard | Medium components |  | pH |  |  | pH and gas flow |
| --- | --- | --- | --- | --- | --- | --- | --- | --- |
| Set-up |  | 1 | 3a | 4 | 2 | 3b | 5 | 6 |
| $Y_{P/S, \text{ used}} \text{ (g/g)}$ | | $0.98 \pm 0.00$ | 0.95 | $0.91 \pm 0.03$ | $0.89 \pm 0.04$ | $0.84 \pm 0.03$ | $0.84 \pm 0.00$ | $0.81 \pm 0.01$ |
| $Y_{P/S, \text{ fed}} \text{ (g/g)}$ | | $0.51 \pm 0.08$ | 0.44 | $0.46 \pm 0.01$ | $0.32 \pm 0.03$ | $0.38 \pm 0.01$ | $0.39 \pm 0.00$ | $0.33 \pm 0.00$ |
| $Y_{P/S, \text{ fixed}} \text{ (g/g)}$ | | $0.99 \pm 0.21$ | 1.00 | $0.93 \pm 0.03$ | $0.93 \pm 0.10$ | $0.94 \pm 0.06$ | $0.93 \pm 0.01$ | $0.92 \pm 0.03$ |
| $Y_{P/X} \text{ (g/g)}$ | | $23.89 \pm 2.58$ | 21.73 | $27.63 \pm 1.44$ | $9.06 \pm 0.62$ | $18.36 \pm 2.33$ | $11.98 \pm 0.33$ | $13.35 \pm 0.56$ |
| $V_{\text{gas, fed}} \text{ (L)}$ | | $78.93 \pm 0.00$ | 73.80 | $76.02 \pm 1.49$ | $25.65 \pm 0.00$ | $73.76 \pm 0.05$ | $49.32 \pm 0.00$ | $53.84 \pm 0.00$ |
| Acetate : Ethanol (mol) | | $13.23 \pm 2.96$ | 11.49 | $11.66 \pm 1.73$ | $9.93 \pm 2.15$ | $11.69 \pm 1.23$ | $6.55 \pm 1.72$ | $5.07 \pm 0.54$ |
| Productivity<br>(g L<sup>-1</sup> h<sup>-1</sup>) | Acetate | $0.25 \pm 0.04$ | 0.22 | $0.23 \pm 0.01$ | $0.17 \pm 0.01$ | $0.19 \pm 0.00$ | $0.19 \pm 0.00$ | $0.13 \pm 0.00$ |
| | EtOH | $0.02 \pm 0.00$ | 0.01 | $0.02 \pm 0.00$ | $0.01 \pm 0.00$ | $0.01 \pm 0.00$ | $0.02 \pm 0.01$ | $0.02 \pm 0.00$ |
| | Total (acetate + ethanol) | $0.27 \pm 0.04$ | 0.23 | $0.25 \pm 0.01$ | $0.18 \pm 0.01$ | $0.20 \pm 0.00$ | $0.21 \pm 0.00$ | $0.15 \pm 0.00$ |

### Effect of pH

When a fermentation with no pH regulation was performed (set-up 2), growth slowed down after 20 h of process-time, with the pH having decreased to 5.0. After 43 h, when the pH value was already at its lowest (4.4), no significant growth or product formation could be detected. An increase in the CDW between 43 and 48 h was observed but was subsequently followed by a further decline and eventually remained mostly constant, with a final value of 0.5 g/L. The final acetate and ethanol concentration achieved were 6.2 and 0.4 g/L, respectively (Figure 3A). Gas consumption stopped after 40 h, with the maximum overall usage interval lasting only 5 h (Figure 3A and Table 2). The yields and productivities for this fermentation where the lowest among all the tests performed, with the exception of Y_P/S, fixed_ (both end-of-process and up to the end of maximum CO fixation) and Y_P/S, used_ (calculated up to the end of the maximum CO fixation), which were analogous to the rest. More detail can be found in tables 3 and 4.

**Figure 3:**
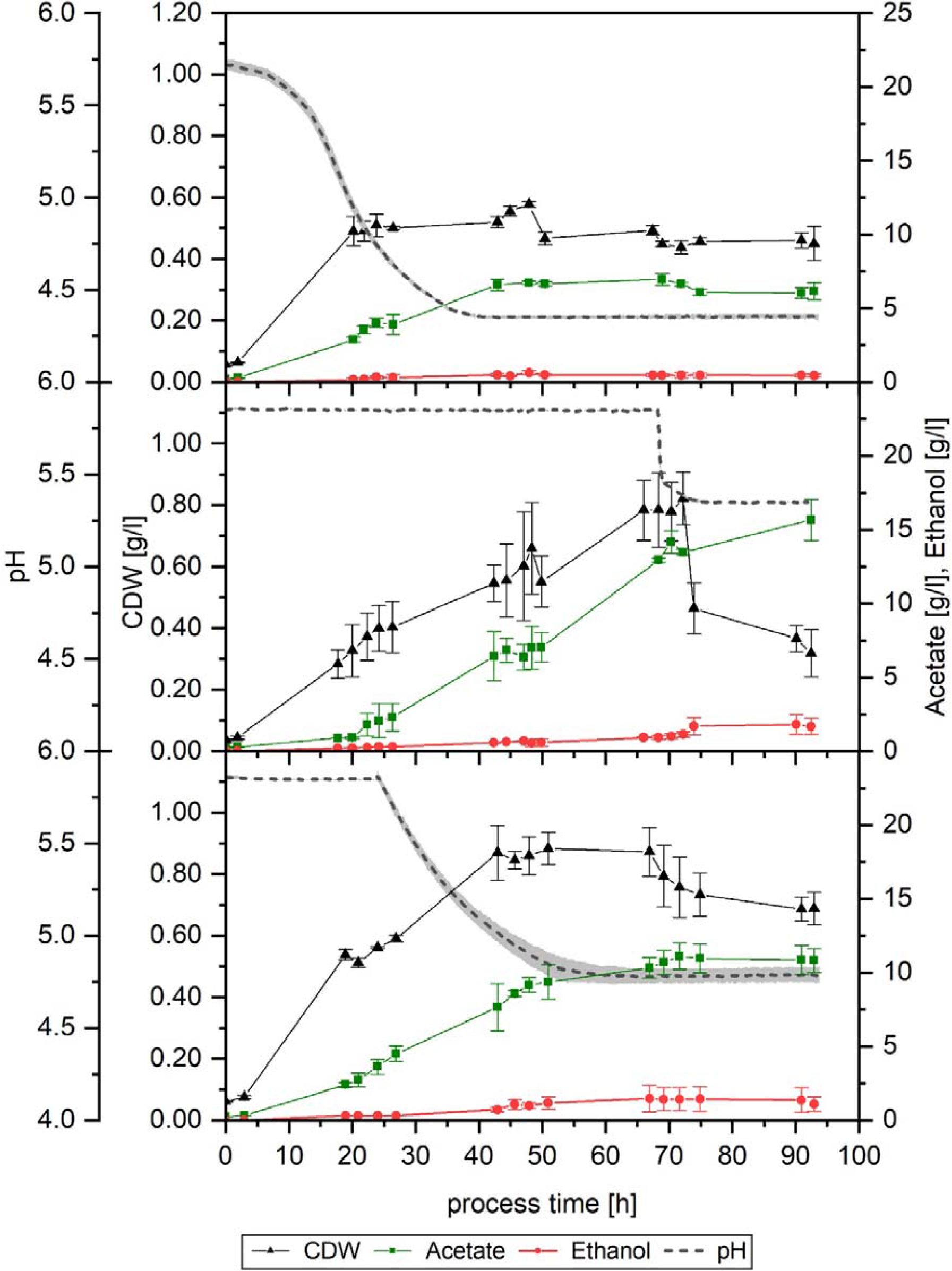
Growth, pH profile and product formation of set-up 2 without pH regulation (A), set-up 3b with increased cysteine and pH change to 5.4 after 68 h (B) and set-up 5 with pH allowed to drop to 4.8 after 24 h (C). Average values of the triplicates (n=3) for cell dry weight (CDW, black triangles), acetate (green squares), ethanol (red dots), and pH (grey dotted line). The lightly colored area around the pH average represent the standard deviation.

In the standard set-up, gas consumption started to decrease after approximately 70 h. After observing that for the first 68 h hours in set-up 3 the gas consumption, growth, and product formation were equivalent to that of set-up 1 (Figures 1A and 1B, and 2A and 2B), the effect of lowering the pH after that point was investigated in set-up 3b. Mainly, the aim was to observe the effect that a lower pH value would have in this late stage of the fermentation, especially regarding the product formation and its ratios. At 68 h, the pH was lowered in two of the fermenters by 0.5 units to 5.4, by using 4 M H_3_PO_4_. As a result, maximum CO fixation came to an end, and a noticeable divergence between fermenters could be noted from this point on. In one fermenter (Figure 4B), immediately after the pH shift the gas consumption started to decrease for CO_2_ and H_2_, and after a small delay, also for CO. Despite the declining tendency, some consumption could be detected up to 92 h: H_2_, CO and CO_2_ average usage was 8, 25 and −16 %, respectively, between 80 and 92 h. CO fixation during this interval was 17 % on average. In the second fermenter (Figure 4C) a drop in H_2_ and CO_2_ usage also happened, but it eventually stabilized at around 50 % and −60 %, respectively. CO usage was still at its maximum, but as a result of the cells not using CO_2_ any further, net CO fixation decreased as well, to an average of 69 % between 68 and 92 h. For the first fermenter, maximum overall usage lasted for 32 h (Table 2, fermenter “c”), while it was 11 h shorter in the second (Table 2, fermenter “d”). Looking at the CDW and product formation (Figure 3B), the deviation between the fermenters is apparent in the biomass yield, as indicated by the standard deviation bars, but much less remarkable in the case of product formation. The maximum CDW measured was, on average, 0.9 g/L at 74 h. After that, the amount of biomass in the fermenter fell to its final value, 0.7 g/L. Acetate was produced throughout the fermentation, even after the biomass decreased. The final concentration obtained was 15.7 g/L. Ethanol, on the other hand, increased until 74 h of process time. Between 72 and 74 h, a somehow steeper increase of 0.4 g/L in the ethanol concentration in the fermenter was detected, from 1.1 to 1.5 g/L, value which remained constant later on. The yields and productivities achieved in this test were, in general, lower compared to the standard, although not to such an extent as seen in the non-pH-regulated fermentation (tables 3 and 4).

**Figure 4:**
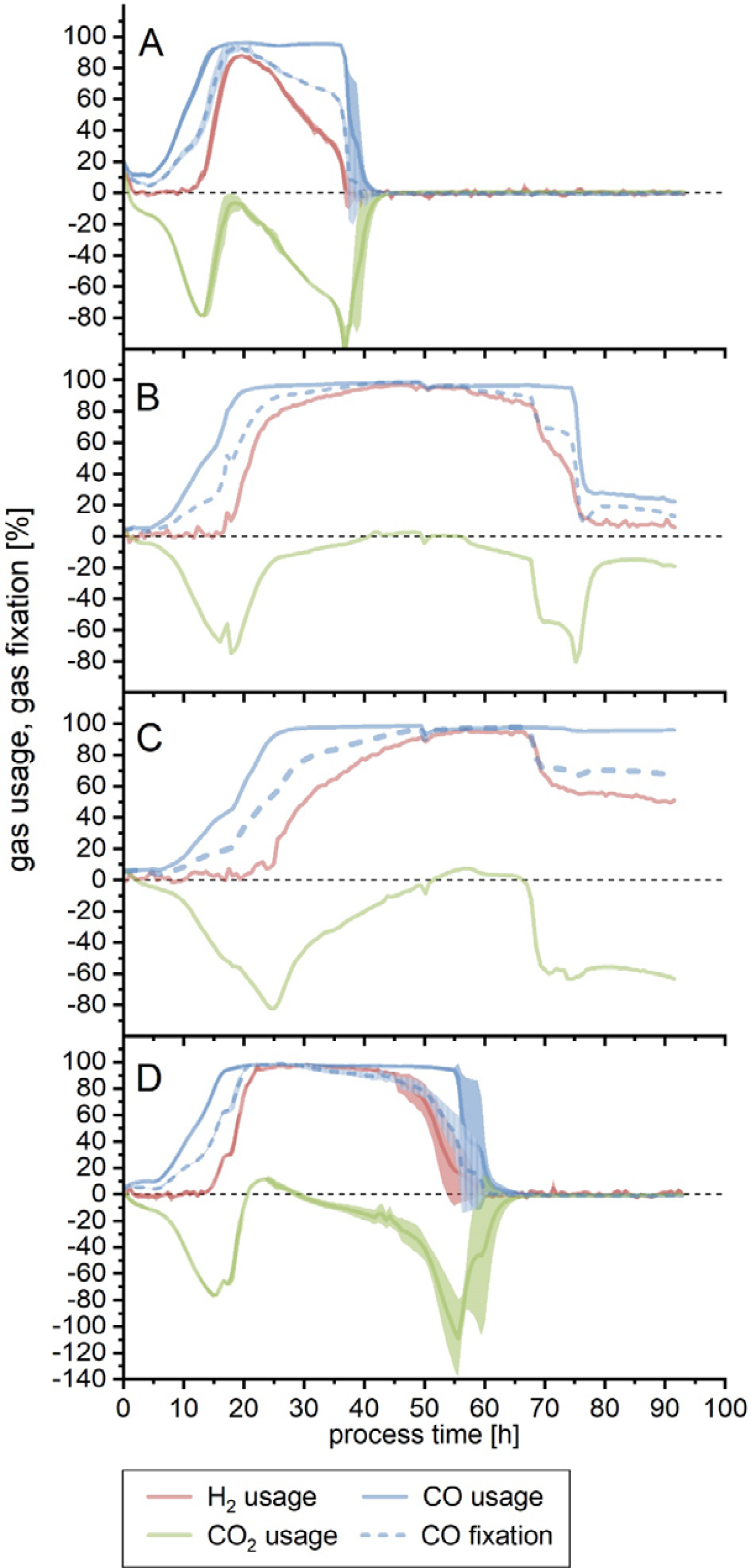
Substrate usage or fixation for set-up 2 without pH regulation (A), set-up 3b with increased cysteine and pH change to 5.4 after 68 h (B and C) and set-up 5 with pH allowed to drop to 4.8 after 24 h (D). Figure B and C show each of the individual fermenter profiles due to the divergence observed between them: the second fermenter (C) is remarkably delayed. Usage is shown for H_2_ (red line), CO_2_ (green line) and CO (blue line). CO fixation is depicted by the dotted blue line. The calculated difference between amount of substance flow rate fed into the bioreactor and the amount of substance flow rate detected in the off-gas is shown here as a percentage. For the CO fixation, if the CO_2_ usage was negative, the amount of CO_2_ produced was subtracted from the amount of (perceived) CO used. Except where otherwise stated, lines show the average of a triplicate (n=3), while the lighter colored areas depict the standard deviation

In the non-pH-regulated fermentation (set-up 2), at 22.5 h just before gas consumption started to diminish, and when the exponential phase had already ended, but there was still cell growth detected, the measured pH was 4.8. This pH value was then chosen for a further test in set-up 5. Here, the fermentation was carried out under standard conditions for 24 h to ensure that gas consumption was already at its highest. As well, in this case, and to avoid a sudden pH change and a potential acid shock in the culture, as well as to avoid the addition of extra salts, the pH was allowed to drop naturally to pH 4.8, and then the pH control was further regulated to this new value, which was reached after 55 h, as Figure 3C shows. In this set-up, biomass concentration reached its maximum earlier than in the standard one: at 43 h, when the pH value was 5.0, the CDW measured was already 0.9 g/L – it was 0.6 g/L in the later. The biomass remained thus stable up to 70 h, dropping after that – 15 h after the pH of 4.8 was reached. Up to 51 h of process-time, acetate formation followed a similar profile to that of set-up 1, reaching a value of 9.4 g/L in the reactor at that time. After this point, though, around the time when the lowest pH was reached and cell concentration decreased, the acetate production slowed down and eventually stopped at 11 g/L, at around 70 h. Ethanol formation also stopped at this point, reaching a final maximum concentration of 1.4 g/L. As can be seen in Figure 4D, during the first 24 h of cultivation the gas consumption followed a trend equivalent to that of the standard set-up, although it reached its maximum 12 h earlier (Table 2). It can also be noted that the maximum overall usage interval was shorter, as well as the time until the end of maximum CO fixation. Yields and productivities for this fermentation were also found to be lower in relation to the standard, and the productivities at the end of the 93 h were almost halved (Table 3).

Finally, looking at the yields and productivities up to the end of the maximum CO fixation phase (Table 4), the non-pH-regulated run achieved again both the lowest Y_P/S, fed_ and Y_P/X_. The values for the standard fermentation were, in all cases, higher than the rest of the set-ups where pH was modified. That being said, in these runs the acetate to ethanol ratio was lower in comparison to the standard, indicating a shift towards more ethanol per mol of acetate produced. Productivities of set-ups 2, 3 (excluding the fermenter where the pH was not changed), and 5 were all similar.

The off-gas profile for set-ups 2, 3b and 5 are shown in Supplementary Figure 2.

### Effect of gas flow

It was noted that in set-up 5, despite the lower pH, the achieved cell growth was similar or even slightly higher than in the standard fermentation, but the product formation was lower. Because of this, the focus was turned to finding out if a reduction in the gas flow, as well as in the pH, would direct the culture towards the formation of more products rather than biomass. In order to do so, set-up 6 was run as set-up 5 for the first 24 h, time after which the pH was allowed to drop naturally until 4.8. At the same time, the gas flow was reduced by 30 % from 18 mL/min to 12.6 mL/min. This flow was deemed adequate to avoid excessive starvation of the culture.

First of all, pH 4.8 was reached at 58 h, 3 h later than in set-up 5, but in this case, and due to the configuration of the pH control, it continued to drop further until 4.7 at 69 h, value at which remained constant thereafter. Concerning cell growth, a CDW of 0.4 g/L was achieved after 24 h, contrasting with the higher CDW of set-up 5 (0.56 g/L), even if the conditions in both runs were equal up to that point. The maximum biomass concentration for this fermentation was also lower: 0.6 at 69 h, coinciding in time with the moment when the pH reached its final lower value. From this point on, no cell growth was detected, and the biomass concentration in the reactor eventually decreased. Acetate was produced until around this time point, as well. Its final concentration, 9 g/L, is lower than in set-up 5 (11 g/L), but not so ethanol: in this last fermentation, 2 g/L could be formed (Figure 5). Looking at the acetate to ethanol ratio, found in tables 3 and 4, this is the fermentation with the lowest value achieved, that is, the product formation is clearly shifted towards ethanol.

**Figure 5:**
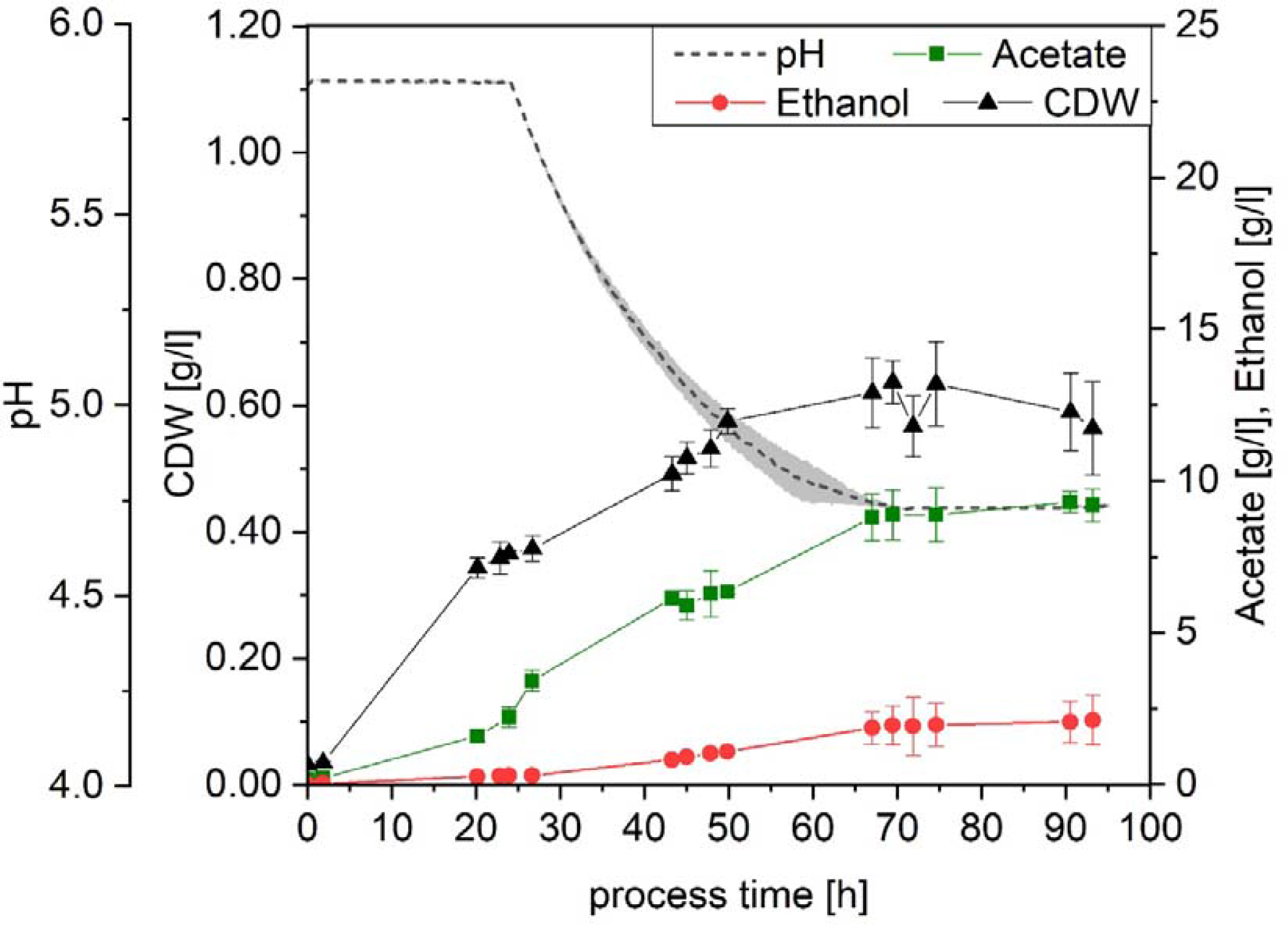
Growth, pH profile and product formation of set-up 6 with pH allowed to drop to 4.8 and gas flow decreased to 12.6 mL/min after 24 h. Average values of the triplicates (n=3) for cell dry weight (CDW, black triangles), acetate (green squares), ethanol (red dots), and pH (grey dotted line). The lightly colored area around the pH average represent the standard deviation.

Gas consumption for the first 24 h was similar in both fermentation 5 and 6 (figures 4D and 6). Maximum gas usage was attained after 24 h in set-up 6, similarly to set-up 5 (21 h) (Table 2). In set-up 6, due to the reduced flow, both the maximum usage interval and the time up to the end of the maximum CO fixation were prolonged (8 and 14 h longer, respectively).

**Figure 6:**
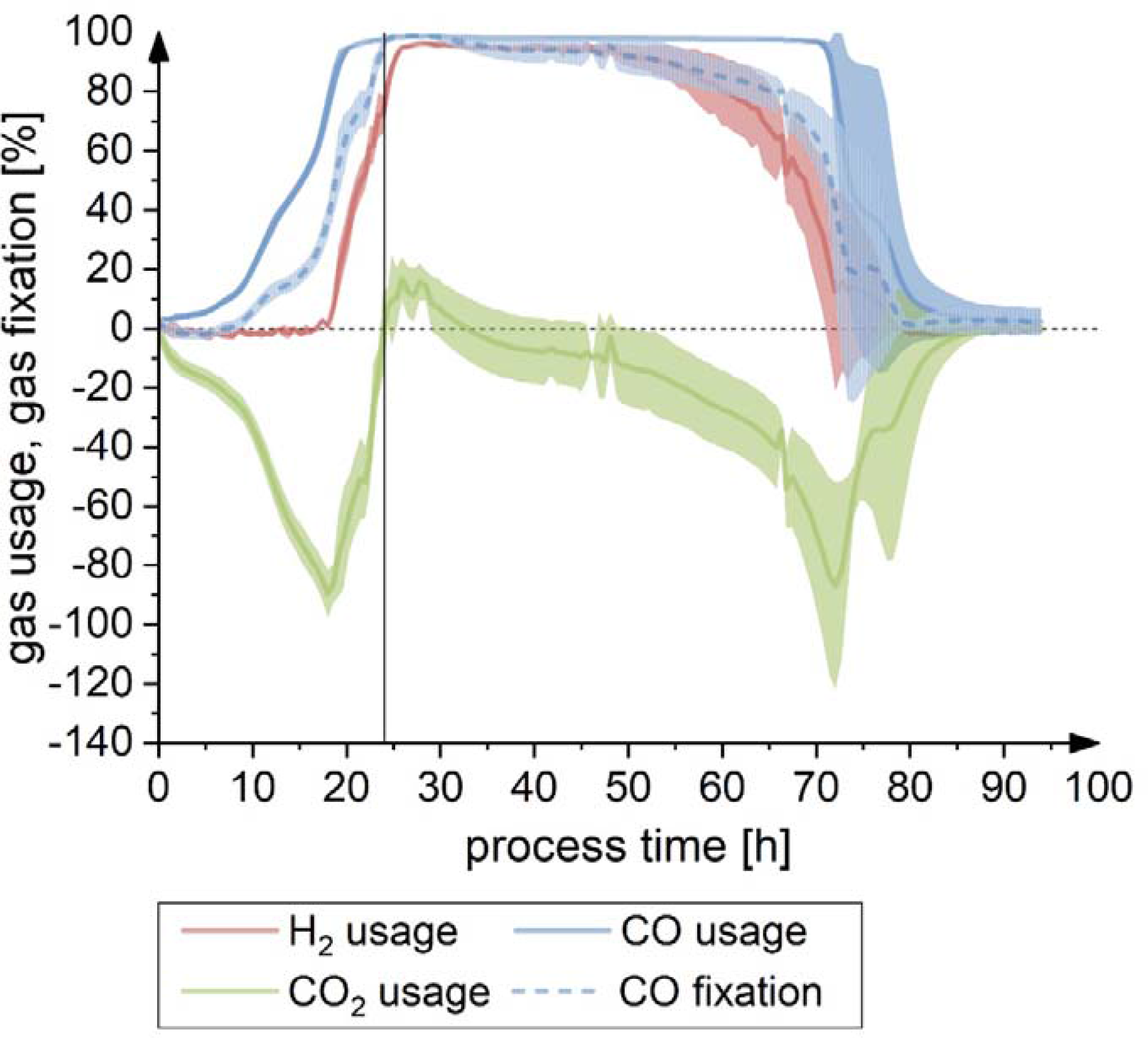
Substrate usage or fixation for set-up 6, where pH was allowed to drop to 4.8 and gas flow was decreased to 12.6 mL/min after 24 h (vertical line). Usage is shown for H_2_ (red line), CO_2_ (green line) and CO (blue line). CO fixation is depicted by the dotted blue line. The calculated difference between amount of substance flow rate fed into the bioreactor and the amount of substance flow rate detected in the off-gas is shown here as a percentage. For the CO fixation, if the CO_2_ usage was negative, the amount of CO_2_ produced was subtracted from the amount of (perceived) CO used. Lines show the average of the triplicate (n=3), while the lighter colored areas depict the standard deviation.

The off-gas profile for set-ups 6 is shown in Supplementary Figure 3.

The yields achieved by lowering the flow after 24 h show that it did not have an impact on how much substrate was fixed into product (Y_P/S, fixed_), given that the results achieved by this fermentation (0.94 ± 0.04 g/g for the complete run, and 0.92 ± 0.03 g/g up to the end of maximum CO consumption) are comparable to the other set-ups (Tables 3 and 4). The product yield per carbon fed was lower than in set-up 5 when calculating it up to the point of the end of maximum CO fixation phase, but contrarily, was improved in this last run in terms of the overall values, as a result of a prolonged gas consumption phase and a lower substrate flow. Due to the diminished growth in set-up 6, the Y_P/X_ calculated at both points was higher, demonstrating that more products had been formed per gram of biomass. Nevertheless, the highest values of the standard set-up could not be reached. The acetate to ethanol ratio, though, was the lowest of the 6 set-ups, being about half of that of set-up 1 (tables 3 and 4): all in all, this fermentation was displaced towards less growth, as well as less acetate and more ethanol per gram of biomass. Despite this, productivities for this set-up where lower than for set-up 5, and they were almost half of those of the standard run.

## DISCUSSION

From the results of the experiments with higher cysteine and yeast extract, it is clear that none of these approaches caused any advantage. It can be excluded that the fact that the culture stopped using the substrates after a certain amount of time was due to a lack of sulfur or nitrogen in the media. In the case of cysteine, a study by Abubackar et al. (2012) with *Clostridium autoethanogenum* reported that an increase in the cysteine-HCl (1.2 g/L vs. 0.5 g/L) had a slightly detrimental impact on biomass, but a higher concentration of ethanol could be reached. The same influence on the biomass was observed on *C. aceticum* with concentrations above 0.5 g/L in another study by Sim and Kamaruddin (2008), although in this case, the product, acetic acid, was not significantly affected. In our case, none of these effects were seen, with cysteine not significantly impacting the fermentation outcome, nor positively or negatively.

The increment of yeast extract in the medium did not translate to higher biomass formation; on the contrary, a slight decrease in biomass could be observed. As well, a higher Y_P/X_ was achieved, and the acetate to ethanol ratio was not significantly altered, especially up to the point when maximum CO fixation stopped. Considering the complete run, only a slight increase in the acetate to ethanol ratio was detected. This is a somewhat unexpected outcome since according to Barik et al. (1988), the higher amount of acetate that they observed with a higher yeast extract concentration was due to enhanced cell growth, which was not the case here. Besides in this study the amount of yeast extract did not have an impact on the final ethanol concentration in our set-up, contrary to what has been reported elsewhere (Abubackar et al., 2012; Barik et al., 1988; Phillips et al., 1993; Vega et al., 1989). One potential argument so as to why yeast extract did not have an effect in our media could be that, since the gas mixture used contained N_2_, *C. ljungdahlii* could potentially be fixing it, as proposed by Richter et al. (2016) and Tremblay et al. (2013), and thus minimizing the effect of other nitrogen sources, but this remains a controversial topic since Emerson et al. (2019) could not observe any nitrogen fixation in their experiments.

Lowering the pH did not achieve the intended effect of shifting the product formation towards ethanol if the whole run is taken into consideration, but looking at the values up to the point when maximum CO consumption ended, all fermentations with a lower pH had an increased amount of ethanol per acetate produced (in mol). Productivities were lower, though, which could have been caused by the less favorable growing conditions. Acetate has been regarded as been a growth-associated product, but different studies disagree on whether ethanol might or might not be growth associated. Barik et al. (1988) and Najafpour and Younesi (2006) report that ethanol is not associated with growth; conversely, Cotter et al. (2009b) showed that *C. ljungdahlii* produced significantly less ethanol when the pH was lowered from 6.8 to 5.5 and concluded that this effect could be related to the slower growth observed. Regarding the experiments here shown, the productivity of ethanol is not constant across the different set-ups, and does not seem to be related to biomass formation; although when the conditions were too detrimental, as in the non-pH regulated run, both product formation and biomass were clearly affected.

The combined effect of lowering the pH and the gas flow was successful in decreasing the cell growth and increasing the ethanol ratio. This agrees with recent research on how ethanol production could be regulated not by gene expression, but by the balance between intracellular and extracellular conditions, that is, total acetate concentration and extracellular pH (Richter et al., 2016). A lower amount of acetate accumulating in the culture broth, as a result of less biomass being formed, would result in less acetate being available intracellularly, and thus the microorganisms could have more time to adapt and shift towards ethanol. At the same time, the lower pH would potentiate this effect, since more undissociated acetic acid could diffuse through the membrane to be available for further conversion into ethanol. Even so, the less favorable conditions of this experiment caused a reduction in the overall productivity.

## CONCLUSION

Neither the supplementation with additional cysteine nor yeast extract increased the duration of the gas consumption, and no dramatic effects on product formation could be observed. Decreasing the pH did not immediately result in higher ethanol formation and impacted the productivity negatively. When, additionally to the pH, the gas flow was also reduced, the reduction in biomass production was significant, as well as a reduction in acetate production and an increased ethanol ratio. Further work will be built upon these results and will look deeper into this and other parameters and their combinations in order to gain a more comprehensive understanding of how the fermentation process here presented can be manipulated.

## Data availability statement

The datasets generated for this study are available on request to the corresponding author.

## Author Contributions

AI conceived and planned the experiments, analyzed the experimental data and drafted the manuscript. AI and MK performed the experiments and collected the experimental data. AN substantially contributed to the conception and design of the experiments. All authors contributed to manuscript revision, read and approved the submitted version.

## Acknowledgements

We acknowledge support by Deutsche Forschungsgemeinschaft and the the KIT-Publication Fund of the Karlsruhe Institute of Technology. We acknowledge the work by Laura Herrmann, who conducted her Bachelor thesis under the supervision of MK.

## REFERENCES

Abubackar, H. N., Veiga, M. C., and Kennes, C. (2012). Biological conversion of carbon monoxide to ethanol: effect of pH, gas pressure, reducing agent and yeast extract. Bioresour. Technol. 114, 518–522. doi:10.1016/j.biortech.2012.03.027.

Ahmed, A., Cateni, B. G., Huhnke, R. L., and Lewis, R. S. (2006). Effects of biomass- generated producer gas constituents on cell growth, product distribution and hydrogenase activity of *Clostridium carboxidivorans* P7^T^. Biomass and Bioenergy 30, 665–672. doi:10.1016/j.biombioe.2006.01.007.

Annan, F. J., Al-Sinawi, B., Humphreys, C. M., Norman, R., Winzer, K., Köpke, M., et al. (2019). Engineering of vitamin prototrophy in *Clostridium ljungdahlii* and *Clostridium autoethanogenum*. Appl. Microbiol. Biotechnol. 103, 4633–4648. doi:10.1007/s00253-019-09763-6.

Barik, S., Prieto, S., Harrison, S. B., Clausen, E. C., and Gaddy, J. L. (1988). Biological production of alcohols from coal through indirect liquefaction. Appl. Biochem. Biotechnol. 18, 363–378. doi:10.1007/BF02930840.

Bengelsdorf, F. R., Beck, M. H., Erz, C., Hoffmeister, S., Karl, M. M., Riegler, P., et al. (2018). Bacterial Anaerobic Synthesis Gas (Syngas) and CO_2_ + H_2_ Fermentation. Adv. Appl. Microbiol. doi:10.1016/bs.aambs.2018.01.002.

Cotter, J. L., Chinn, M. S., and Grunden, A. M. (2009a). Ethanol and acetate production by *Clostridium ljungdahlii* and *Clostridium autoethanogenum* using resting cells. Bioprocess Biosyst. Eng. 32, 369–380. doi:10.1007/s00449-008-0256-y.

Cotter, J. L., Chinn, M. S., and Grunden, A. M. (2009b). Influence of process parameters on growth of *Clostridium ljungdahlii* and *Clostridium autoethanogenum* on synthesis gas. Enzyme Microb. Technol. 44, 281–288. doi:10.1016/j.enzmictec.2008.11.002.

De Tissera, S., Köpke, M., Simpson, S. D., Humphreys, C., Minton, N. P., and Dürre, P. (2017). “Syngas biorefinery and syngas utilization,” in Biorefineries. Advances in Biochemical Engineering/Biotechnology, eds. K. Wagemann and N. Tippkötter (Springer, Cham), 247–280. doi:10.1007/10_2017_5.

Diender, M., Stams, A. J. M., and Sousa, D. Z. (2016). Production of medium-chain fatty acids and higher alcohols by a synthetic co-culture grown on carbon monoxide or syngas. Biotechnol. Biofuels 9, 82. doi:10.1186/s13068-016-0495-0.

Edenhofer, O., Pichs-Madruga, R., Sokona, Y., Kadner, S., Minx, J. C., Brunner, S., et al. (2014). “Technical Summary,” in Climate Change 2014: Mitigation of Climate Change. Contribution of Working Group III to the Fifth Assessment Report of the Intergovernmental Panel on Climate Change, eds. O. Edenhofer, R. Pichs-Madruga, Y. Sokona, E. Farahani, S. Kadner, K. Seyboth, et al. (Cambridge University Press, Cambridge, United Kingdom and New York, NY, USA), 31–107. doi:10.1017/CBO9781107415416.

Emerson, D. F., Woolston, B. M., Liu, N., Donnelly, M., Currie, D. H., and Stephanopoulos, G. (2019). Enhancing hydrogen-dependent growth of and carbon dioxide fixation by *Clostridium ljungdahlii* through nitrate supplementation. Biotechnol. Bioeng. 116, 294–306. doi:10.1002/bit.26847.

Henstra, A. M., Sipma, J., Rinzema, A., and Stams, A. J. (2007). Microbiology of synthesis gas fermentation for biofuel production. Curr. Opin. Biotechnol. 18, 200–206. doi:10.1016/j.copbio.2007.03.008.

Hu, P., Chakraborty, S., Kumar, A., Woolston, B., Liu, H., Emerson, D., et al. (2016). Integrated bioprocess for conversion of gaseous substrates to liquids. Proc. Natl. Acad. Sci. 113, 3773–3778. doi:10.1073/pnas.1516867113.

Klasson, K. T., Ackerson, M. D., Clausen, E. C., and Gaddy, J. L. (1993). Biological conversion of coal and coal-derived synthesis gas. Fuel 72, 1673–1678. doi:10.1016/0016-2361(93)90354-5.

Kundiyana, D. K., Wilkins, M. R., Maddipati, P., and Huhnke, R. L. (2011). Effect of temperature, pH and buffer presence on ethanol production from synthesis gas by “Clostridium ragsdalei”. Bioresour. Technol. 102, 5794–5799. doi:10.1016/j.biortech.2011.02.032.

Lee, S. Y., Park, J. H., Jang, S. H., Nielsen, L. K., Kim, J., and Jung, K. S. (2008). Fermentative Butanol Production by Clostridia. Biotechnol. Bioeng. 101, 209–228. doi:10.1002/bit.22003.

Liew, F. M. F. F. M., Martin, M. E., Tappel, R. C. R. R. C., Heijstra, B. D. B. B. D., Mihalcea, C., Köpke, M., et al. (2016). Gas Fermentation – A Flexible Platform for Commercial Scale Production of Low Carbon Fuels and Chemicals from Waste and Renewable Feedstocks. Front. Microbiol. 7:694, 1–28. doi:10.3389/fmicb.2016.00694.

Liew, F. M., Köpke, M., and Simpson, S. D. (2013). “Gas Fermentation for Commercial Biofuels Production,” in Liquid, Gaseous and Solid Biofuels - Conversion Techniques, ed. Z. Fang (IntechOpen), 125–173. doi:10.5772/52164.

Mohammadi, M., Najafpour, G. D., Younesi, H., Lahijani, P., Uzir, M. H., and Mohamed, A. R. (2011). Bioconversion of synthesis gas to second generation biofuels: A review. Renew. Sustain. Energy Rev. 15, 4255–4273. doi:10.1016/j.rser.2011.07.124.

Najafpour, G., and Younesi, H. (2006). Ethanol and acetate synthesis from waste gas using batch culture of *Clostridium ljungdahlii*. Enzyme Microb. Technol. 38, 223–228. doi:10.1016/j.enzmictec.2005.06.008.

Oswald, F., Dörsam, S., Veith, N., Zwick, M., Neumann, A., Ochsenreither, K., et al. (2016). Sequential Mixed Cultures: From Syngas to Malic Acid. Front. Microbiol. 7, 1–12. doi:10.3389/fmicb.2016.00891.

Phillips, J. R., Klasson, K. T., Clausen, E. C., and Gaddy, J. L. (1993). Biological production of ethanol from coal synthesis gas - Medium development studies. Appl. Biochem. Biotechnol. 39, 559–571. doi:10.1007/BF02919018.

Richter, H., Molitor, B., Wei, H., Chen, W., Aristilde, L., and Angenent, L. T. (2016). Ethanol production in syngas-fermenting *Clostridium ljungdahlii* is controlled by thermodynamics rather than by enzyme expression. Energy Environ. Sci. 9, 2392–2399. doi:10.1039/C6EE01108J.

Saxena, J., and Tanner, R. S. (2012). Optimization of a corn steep medium for production of ethanol from synthesis gas fermentation by *Clostridium ragsdalei*. World J. Microbiol. Biotechnol. 28, 1553–1561. doi:10.1007/s11274-011-0959-0.

Sim, J. H., and Kamaruddin, A. H. (2008). Optimization of acetic acid production from synthesis gas by chemolithotrophic bacterium – *Clostridium aceticum* using statistical approach. Bioresour. Technol. 99, 2724–2735. doi:10.1016/j.biortech.2007.07.004.

Tanner, R. S. (2007). “Cultivation of Bacteria and Fungi,” in Manual of Environmental Microbiology, Third Edition, eds. C. Hurst, R. Crawford, J. Garland, D. Lipson, A. Mills, and L. Stetzenbach (ASM Press, Washington, DC), 69–78. doi:10.1128/9781555815882.ch6.

Tremblay, P.-L., Zhang, T., Dar, S. A., Leang, C., and Lovley, D. R. (2013). The Rnf Complex of *Clostridium ljungdahlii* Is a Proton-Translocating Ferredoxin:NAD^+^ Oxidoreductase Essential for Autotrophic Growth. Appl. Environ. Microbiol. 8, 1–8. doi:10.1128/mBio.00406-12.Editor.

Vega, J. L., Prieto, S., Elmore, B. B., Clausen, E. C., and Gaddy, J. L. (1989). The Biological Production of Ethanol from Synthesis Gas. Appl. Biochem. Biotechnol. 20–21, 781–797. doi:10.1007/BF02936525.

Worden, R. M., Grethlein, A. J., Jain, M. K., and Datta, R. (1991). Production of butanol and ethanol from synthesis gas via fermentation. Fuel 70, 615–619. doi:10.1016/0016-2361(91)90175-A.

