## Supplementary figures and images for "Effect of Cysteine, Yeast Extract, pH Regulation and Gas Flow on Acetate and Ethanol Formation and Growth Profiles of *Clostridium ljungdahlii* Syngas Fermentation"

### Supplemental Figure 1

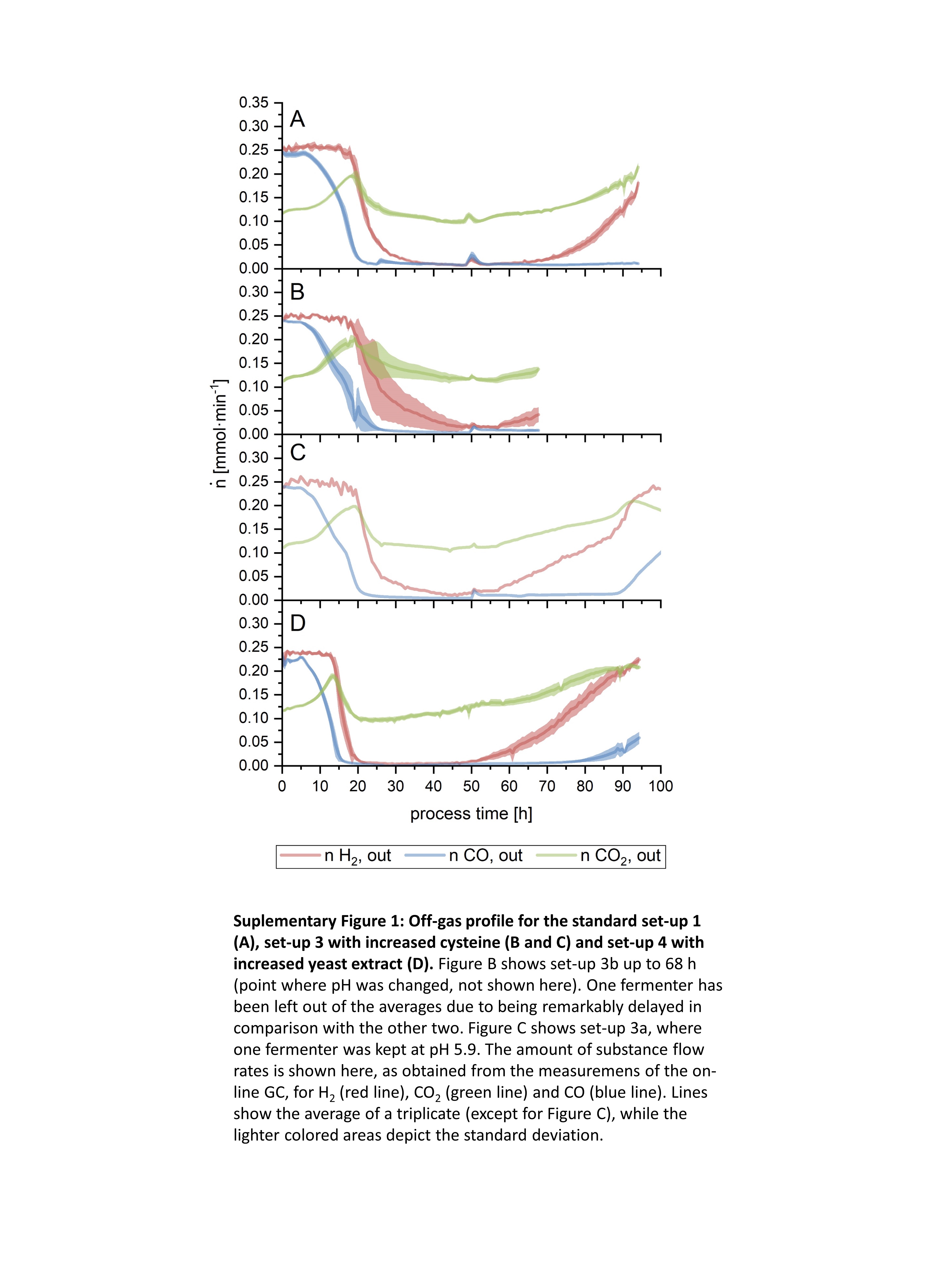

### Supplemental Figure 2

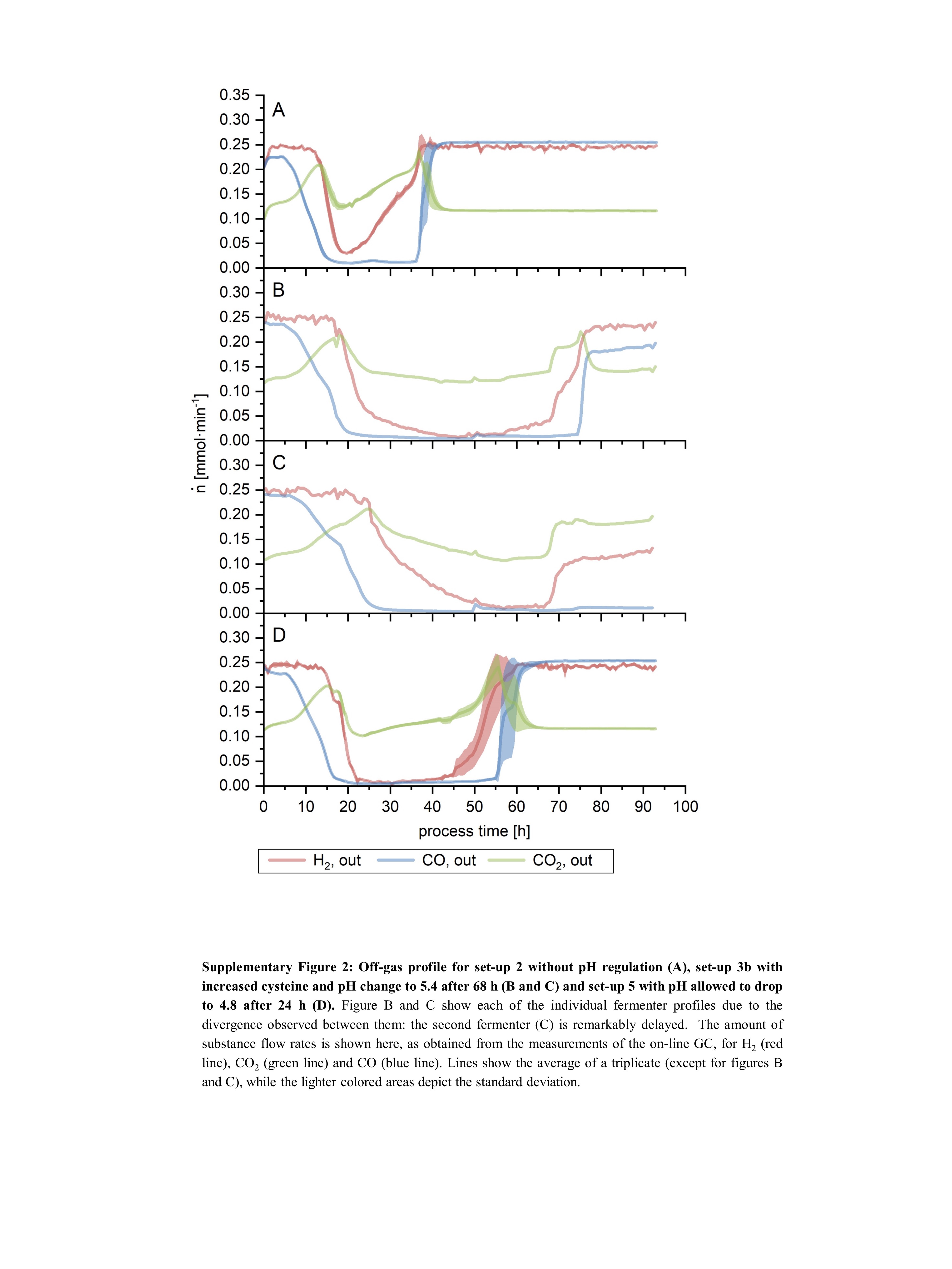

### Supplemental Figure 3

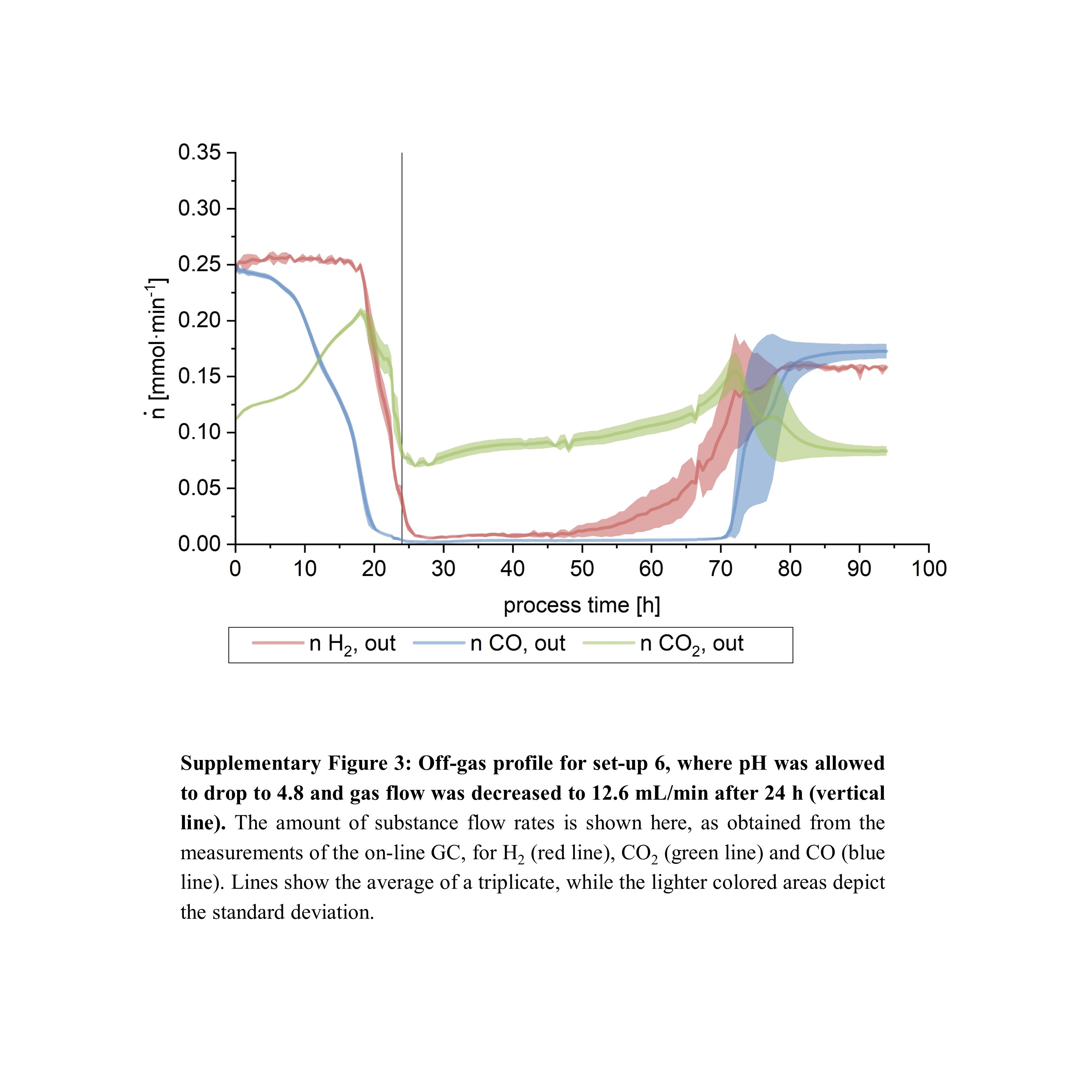
